# Enabling out-of-clinic human immunity studies via single-cell profiling of capillary blood

**DOI:** 10.1101/2020.07.25.210468

**Authors:** Tatyana Dobreva, David Brown, Jong Hwee Park, Matt Thomson

**Author notes:** These authors contributed equally (Tatyana Dobreva, David Brown).

## Abstract

An individual’s immune system is driven by both genetic and environmental factors that vary over time. To better understand the temporal and inter-individual variability of gene expression within distinct immune cell types, we developed a platform that leverages multiplexed single-cell sequencing and out-of-clinic capillary blood extraction to enable simplified, cost-effective profiling of the human immune system across people and time at single-cell resolution. Using the platform, we detect widespread differences in cell type-specific gene expression between subjects that are stable over multiple days.

**Summary:** Increasing evidence implicates the immune system in an overwhelming number of diseases, and distinct cell types play specific roles in their pathogenesis.^1,2^ Studies of peripheral blood have uncovered a wealth of associations between gene expression, environmental factors, disease risk, and therapeutic efficacy.^4^ For example, in rheumatoid arthritis, multiple mechanistic paths have been found that lead to disease, and gene expression of specific immune cell types can be used as a predictor of therapeutic non-response.^12^ Furthermore, vaccines, drugs, and chemotherapy have been shown to yield different efficacy based on time of administration, and such findings have been linked to the time-dependence of gene expression in downstream pathways.^21,22,23^ However, human immune studies of gene expression between individuals and across time remain limited to a few cell types or time points per subject, constraining our understanding of how networks of heterogeneous cells making up each individual’s immune system respond to adverse events and change over time.

## Introduction

The advent of single-cell RNA sequencing (scRNA-seq) has enabled the interrogation of heterogeneous cell populations in blood without cell type isolation and has already been employed in the study of myriad immune-related diseases.^14,15,18^ Recent studies employing scRNA-seq to study the role of immune cell subpopulations between healthy and ill patients, such as those for Crohn’s disease ^41^, Tuberculosis ^40^, and COVID-19 ^39^, have identified cell type-specific disease relevant signatures in peripheral blood immune cells; however, these types of studies have been limited to large volume venous blood draws which can tax already ill patients, reduce the scope of studies to populations amenable to blood draws, and often require larger research teams to handle the patient logistics and sample processing costs and labor. In particular, getting repeated venous blood draws within a single day and/or multiple days at the subject’s home has been a challenge for older people with frail skin and those on low dosage Acetylsalicylic acid.^44^ This dependence on venous blood dramatically impacts our ability to understand the high temporal dynamics of health and disease.

Capillary blood sampling is being increasingly used in point-of-care testing and has been advised for obese, elderly, and other patients with fragile or inaccessible veins.^20,42,43,15^ The reduction of patient burden via capillary blood sampling could enable researchers to perform studies on otherwise difficult or inaccessible populations, and at greater temporal resolution. To date, scRNA-seq of human capillary blood has not yet been validated nor applied to study the immune system. In order to make small volumes of capillary blood (100 ul) amenable to scRNA-seq we have developed a platform which consists of a painless vacuum-based blood collection device, sample de-multiplexing leveraging commercial genotype data, and an analysis pipeline used to identify time-of-day and subject specific genes. The potential of our platform is rooted in enabling large scale studies of immune state variation in health and disease across people. The high-dimensional temporal transcriptome data could be paired with computational approaches to predict and understand emergence of pathological immune states. Most importantly, our platform makes collection and profiling of human immune cells less invasive, less expensive and as such more scalable than traditional methods rooted in large venous blood draws.

## Results

### Platform for low-cost interrogation of single-cell immune gene expression profiles

Our platform is comprised of a protocol for isolating capillary peripheral blood mononuclear cells (CPBMCs) using a push-button collection device (TAP)^15^, pooling samples to reduce per-sample cost using genome-based demultiplexing^16^, and a computational package that leverages repeated sampling to identify genes that are differentially expressed in individuals or between time points, within subpopulations of cells (Fig. 1a). Using a painless vacuum-based blood collection device such as the commercial FDA-approved TAP to collect capillary blood makes it convenient to perform at-home self-collected sampling and removes the need for a trained phlebotomist, increasing the ease of acquiring more samples. The isolation of CPBMCs is done using gradient centrifugation and red blood cells are further removed via a red blood cell lysis buffer. The cells from the different subjects are pooled, sequenced via scRNA-seq using a single reagent kit, and de-multiplexed ^16^ via each subject’s single-nucleotide polymorphisms (SNPs), reducing the per-sample processing cost. Finally, we made our pipeline compatible with genotyping data obtained from a commercial service (23AndMe), removing the need for a separate genotyping assay for subjects that already have access to their own SNP data.

**Fig.1.**
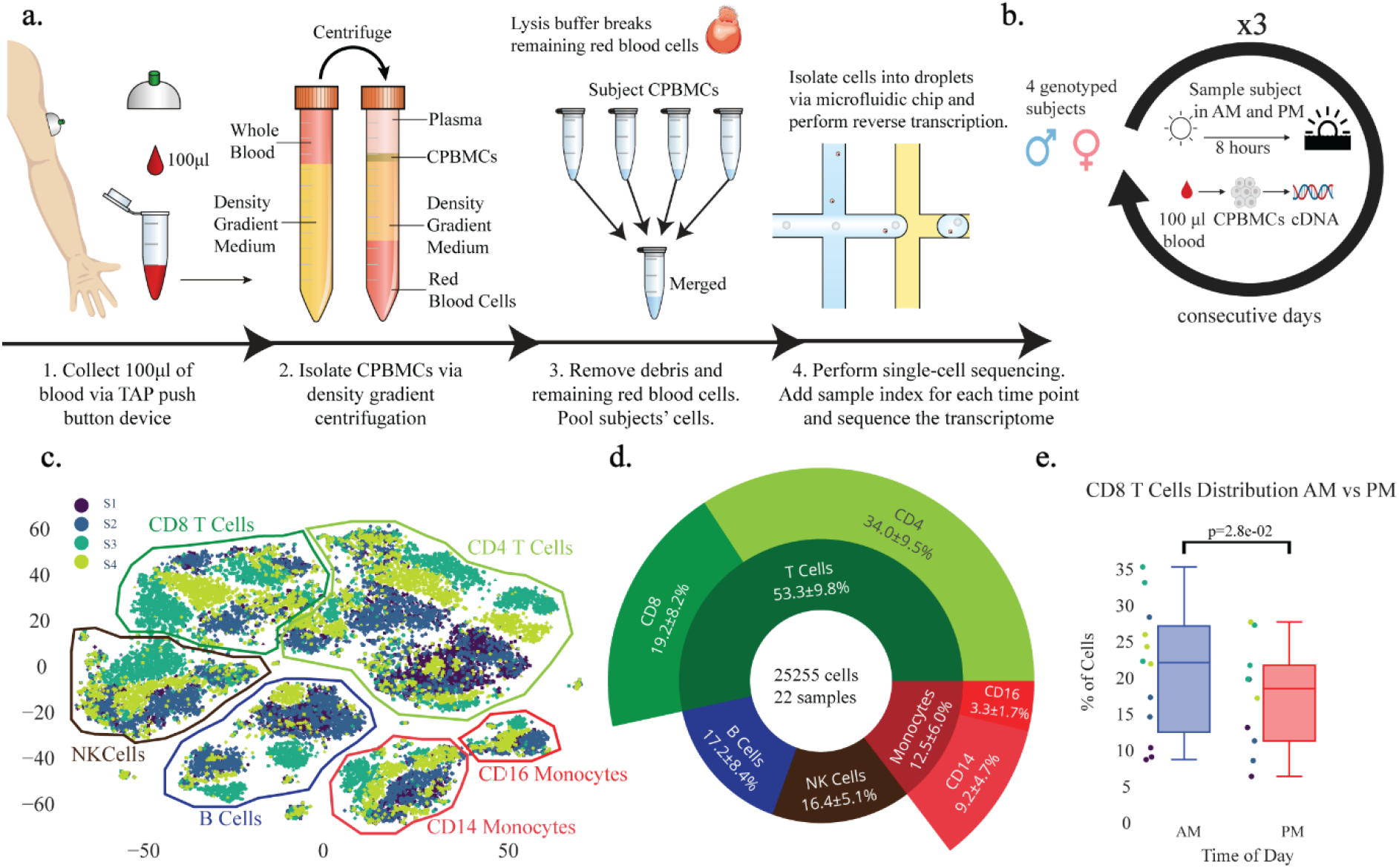
Experimental workflow and consistency of capillary blood sampling. **(a)** Experimental workflow for capillary blood immune profiling 1. Blood is collected using the TAP device from the deltoid. 2. Capillary peripheral blood mononuclear cells (CPBMCs) are separated via centrifugation. 3. Red blood cells are lysed and removed, and samples from different subjects are pooled together. 4. Cell transcriptomes are sequenced using single-cell sequencing. **(b)** Time-course study design. CPBMCs are collected and profiled from 4 subjects (2 male, 2 female) each morning (AM) and afternoon (PM) for 3 consecutive days. **(c)** 2-dimensional t-SNE projection of the transcriptomes of all cells in all samples. Cells appear to cluster by major cell type, and then further by subject. **(d)** Immune cell type percentages across all samples shows stable cell type abundances (includes cells without subject labels). **(e)** CD8^+^ T cells have significantly lower abundance in PM samples vs AM samples, consistent with previous findings from venous blood (student t-test performed on AM, PM samples of each subject and then combined using Stouffer’s method). Marker colors correspond to subject identity from **(c)**.

### Single-cell RNA sequencing (scRNA-seq) of low volume capillary blood recovers distinct immune cell populations stably across time

As a proof-of-concept, we leveraged our scRNA-seq of capillary blood platform to identify genes that exhibit diurnal behavior in subpopulations of cells and find subject-specific immune relevant gene signatures. We performed a three-day study in which we processed capillary blood from four subjects in the morning and afternoon, totaling 25,255 cells across 22 samples (Fig. 1b). Major immune cell types such as T cells (CD4^+^, CD8^+^), Natural Killer cells, Monocytes (CD14^+^, CD16^+^), and B cells are present in all subjects and time points with stable expression of key marker genes (Fig. 1d, Fig. S1), demonstrating that these signals are robust to technical and biological variability of CPBMC sampling. Furthermore, cells within subjects cluster closer together than cells between subjects, suggesting that subjects have unique transcriptomic fingerprints (Fig. 1c). Additionally, we observed significantly higher (p=2.8×10^−2^, one-sided student t-test) CD8^+^ T cell abundance in the morning across all subjects (Fig. 1e), corroborating previous findings from venous PBMCs about the daytime dependence of the adaptive immune system composition.^7^

### High frequency scRNA-seq unveils new diurnal cell type-specific genes

Genes driven by time-of-day expression, such as those involved in leukocyte recruitment^27^ and regulation of oxidative stress^28^, have been determined to play an important role in both innate and adaptive immune cells^8^. Medical conditions such as atherosclerosis, parasite infection, sepsis, and allergies display distinct time-of-day immune responses in leukocytes^29^, suggesting the presence of diurnally expressing genes that could be candidates for optimizing therapeutic efficacy via time-of-day dependent administration. However, studies examining diurnal gene expression in human blood have been limited to whole blood gene panels via qPCR, or bulk RNA-seq.^9,37, 38^

Leveraging our platform, which enables single-cell studies of temporal human immune gene expression, we detected 366 genes (FDR < 0.05, multiple comparison corrected) exhibiting diurnal activity within at least one cell subpopulation (Fig. 2a). Among the 20 top diurnally classified genes, we found that 50% of those genes were previously correlated with circadian behavior (Table S1), such as DDIT4^9^ (Fig. 2b), SMAP2^26^, and PDIA3^36^. However, only 101/366 (27.6%) of these genes are detected as diurnal at the whole population level (FDR < 0.05, multiple comparison corrected), suggesting there may be many more diurnally-varying genes than previously discovered. For example, IFI16 and ARID1A (Fig. 2c) have diurnal expression only in NK cells and B cells, respectively, and display previously unreported transcriptional diurnal patterns. In particular, ARID1A is a regulator of higher order chromatin structure and mutations of this gene have been implicated in B-cell lymphomas.^18^ Given previous evidence of increased efficacy of time-dependent chemotherapy administration^23,24^ and tumor cells exhibiting out-of-sync behavior compared to normal cells^25^, understanding ARID1A’s diurnal expression pattern can potentially guide timely administration of candidate therapeutics. Out of the identified 366 diurnally-varying genes, 162 (44%) are considered druggable under the drug gene interaction database (http://www.dgidb.org/).

**Fig.2.**
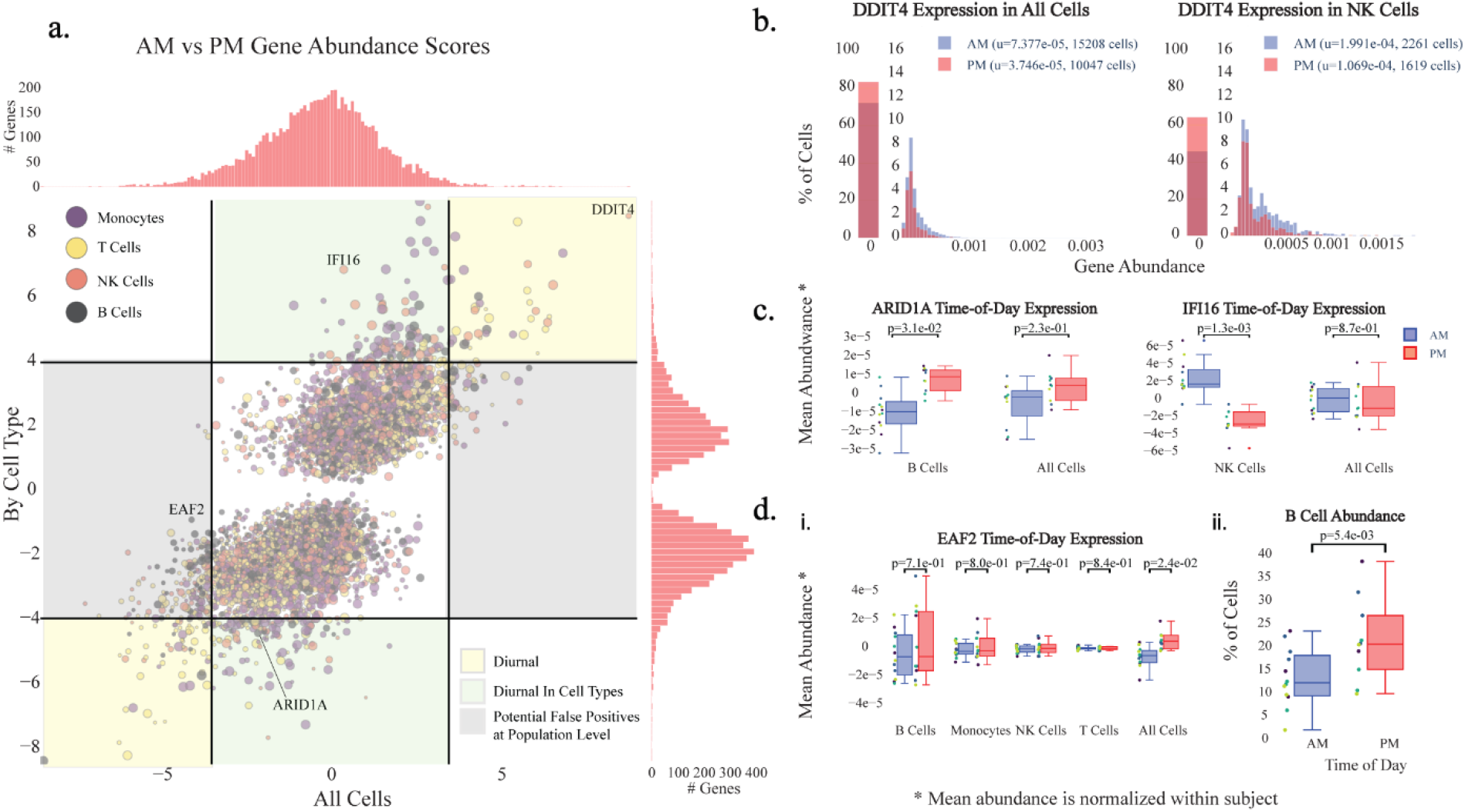
Diurnal variability in subpopulations of capillary blood. **(a)** Magnitude (Z-score) of the difference in AM vs PM gene expression across the whole population of cells (x) vs the cell type with the largest magnitude Z-score (y). Points above or below the significance lines (FDR < 0.05, multiple comparison correction) display different degrees of diurnality. The size of each marker indicates the abundance of the gene (the largest percent of cells in a subpopulation that express this gene). **(b)** Distribution of expression of DDIT4, a previously identified circadian rhythm gene^9^, shows diurnal signal across all cells, as well as individual cell types, such as natural killer (NK) cells. u indicates the mean fraction of transcripts per cell (gene abundance). **(c)** Example of newly identified diurnal genes, ARID1A and IFI16 that could be missed if analyzed at the population level **(d)** Example of a gene, EAF2, that could be falsely classified as diurnal **(i)** without considering cell type subpopulations due to a diurnal B cell abundance shift **(ii)**.

### scRNA-seq profiling distinguishes diurnal gene expression from cell type abundance changes

We also detected 250 genes (FDR < 0.05, multiple comparison corrected) exhibiting diurnal behavior when analyzed at the population level, such as EAF2, that do not display diurnal variation in any of our major cell types (Fig. 2d.i). Such false positives may come from diurnal shifts in cell type abundance rather than up- or down-regulation of genes. In the case of EAF2, which is most abundant in B cells, we hypothesized that the diurnality detected at the population level was a result of an increase of B cell abundance in the afternoon, and verified this in our data (p=5.0×10^−3^, one-sided student-t test) (Fig. 2d.ii). This finding highlights the importance of looking at expression within multiple cell types to avoid potentially misleading mechanistic hypotheses.

### Individuals exhibit robust cell type-specific differences in genes and pathways relevant to immune function

Gene expression studies of isolated cell subpopulations across large cohorts of people have revealed a high degree of variability between individuals that cannot be accounted for by genetics alone, with environmental effects that vary over time likely playing a critical role.^33, 35^ Furthermore, these transcriptomic differences have been linked to a wide range of therapeutic responses, such as drug-induced cardiotoxicity.^34^ However, while immune system composition and expression has been shown to be stable over long time periods within an individual, acute immune responses generate dramatic immune system changes, meaning that large single time point population studies are unable to establish whether variability between individuals is stable or the result of dynamic response to stimuli.^32^

To probe the stability of individual gene expression signatures at the single-cell level, we used our pipeline to identify genes whose variation in gene expression is most likely caused by intrinsic intersubject differences rather than high frequency immune system variability. We compared the mean gene expressions of all time points between subjects in all cell types and identified 1251 genes (FDR < 0.05, multiple comparison corrected) that are differentially expressed in at least one subpopulation of cells. Like Whitney, et al., we found MHC class II genes, such as HLA-DRB1, HLA-DQA2, and HLA-C (Fig. 3a) to be among the largest sources of variation between subjects.^5^ Additionally, we found that DDX17, which was classified by Whitney et al. as a gene with high intersubject variability, but low intrasubject variability via repeat sampling over longer time scales, may be a new class of temporally varying gene that varies by day of week, having consistently increasing expression each subsequent sampling day. This stresses the importance of high frequency sampling for identifying genes with the most intrinsic interindividual variability.

**Fig.3.**
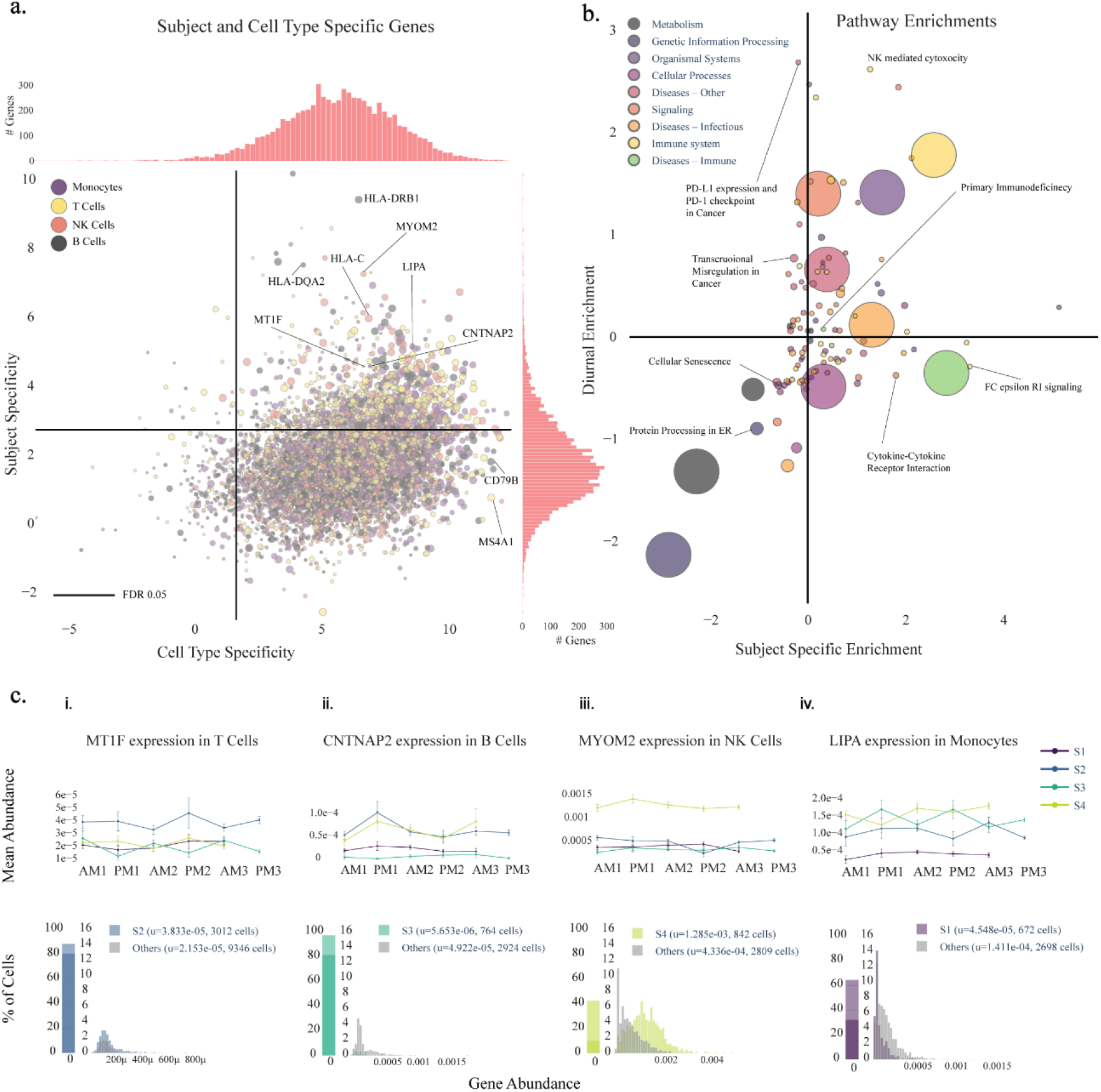
Subject variability in immune and disease-relevant genes and pathways. **(a)** Magnitude (log2 F statistic) of the variability in expression of genes between different cell types (x) and between subjects (y). 1251/7034 (17.8%) of genes are above the subject specificity significance line (FDR < 0.05, multiple comparison correction) and are classified as subject-specific. Several MHC class II genes (HLA-X) are strongly subject-specific, consistent with previous findings^5^. **(b)** KEGG pathways grouped into categories and their enrichment (Z-score from 2-proportion Z-test) among the top 250 diurnally and subject-varying genes vs all genes. Immune system and disease pathways are significantly enriched (p=0.059), supportive of the conclusion that immune and disease-related genes are highly subject dependent. The large circles indicate the enrichment of the category overall, and the sizes of the smaller pathway points indicate the number of genes associated with the pathway. **(c)** Subject and cell type specific gene examples for each subject and cell type with the upper row displaying the trace of mean gene expression across time-points and the bottom row showing gene abundance shifts for the subjects of interest.

### Numerous subject-specific genes are revealed in specific immune cell types

Within the 1251 genes with intrinsic interindividual variability, we found myriad disease-relevant genes for all subjects and cell types, which can be explored at our interactive online portal (http://capblood-seq.caltech.edu). As just one example, subject S1’s monocytes have a consistent downregulation (p=2.0×10^−6^, two-sided student t-test) of LIPA, a gene that is implicated in Lysosomal Acid Lipase Deficiency (Fig. 3c). Given the low abundance of monocytes in blood samples, such findings would typically only be discovered from a targeted blood test or RNA sequencing of isolated monocytes, either of which would only be performed if the disease was already suspected; this showcases how automated discovery in heterogeneous cell populations can be leveraged for personalized, preventative care.

### Immune function and disease pathways are enriched in subject-specific genes

Given that genes do not act alone, we also found cell type-specific pathway differences among subjects. In particular, Subject 2’s S100A8, S100A9, and S100A12 genes, calcium-binding proteins that play an important role in macrophage inflammation, are significantly downregulated in monocytes (p_S100A8_=3.1×10^−5^, p_S100A9_=1.4×10^−4^, p_S100A12_=5.1×10^−4^, two-sided student t-test) compared to other subjects (Fig. S2). We further explored our findings by inspecting the pathways that are most enriched in individual and time-varying genes, and found that genes that are implicated in immune system function (p=0.059) and immune diseases (p=0.041) are more present in subject-specific genes (Fig. 3b). This stands in contrast to pathways of core cellular functions such as genetic information processing (p=0.041) and metabolism (p=0.105), which are less present in subject-specific genes.

## Discussion

Genome and transcriptome sequencing projects have unveiled millions of genetic variants and associated gene expression traits in humans.^10^ However, large-scale studies of their functional effects performed through venous blood draws require tremendous effort to undertake, and this is exacerbated by the cost and complexity of single-cell transcriptome sequencing. Efforts such as the Immune Cell Census^13^ are already underway to perform single-cell profiling of large cohorts, but reliance on venous blood draws of PBMCs will likely limit the diversity and temporal resolution of their sample pool. Our platform gives researchers direct, scalable access to high resolution immune system transcriptome information of human subjects, lowering the barrier of entry for myriad new research avenues. Examples of such studies include: 1. tracking vulnerable populations over time, such as monitoring clonal expansion of CD8^+^ T cells in Alzheimer’s disease progression^2^, 2. profiling of individuals who are under home care to track disease progression and therapeutic response, such as transplant patients and people under quarantine, and 3. tracking how stress, diet, and environmental conditions impact the immune system at short and long time scales, particularly in underrepresented populations who do not have easy access to hospitals or research institutions, such as people in rural or underdeveloped areas. Larger, more diverse subject pools coupled with time course studies of cell type gene expression in both health and disease will have a dramatic impact on our ability to understand the baseline and variability of immune function and disease response.

## Online Content

Online web portal is available to explore data presented in the main figures for study summary, diurnal and subject specific genes via http://capblood-seq.caltech.edu.

## Data Availability

Gene expression matrix and relevant metadata are available on https://data.caltech.edu/records/1407. Genome and FASTQ files are not being released to protect the identity of the subjects.

## Code Availability

Custom code made for diurnal and subject specific gene detection is available on https://github.com/thomsonlab/capblood-seq

## Methods

### Human Study Cohort

This study was conducted at Caltech. Four healthy adults (2 male, 2 female) were recruited. All participants provided written informed consent. The study was approved by the Institutional Review Board (IRB) at Caltech and all methods were performed in compliance with relevant guidelines and regulations. The blood collection took place in a non-BSL room to make sure the subjects were not exposed to pathogens. Subject blood was collected roughly 8 hours apart over three consecutive days.

### CPBMC isolation

100 μl of capillary blood was collected via push-button collection device (TAP from Seventh Sense Biosystems). For each blood draw, the site of collection was disinfected with an alcohol wipe and the TAP device was placed on the deltoid of the subject per device usage instructions. The button was pushed, and then blood was collected for 2-7 minutes until the indicator turned red. Blood was extracted from the TAP device by gently breaking the seal foil, and mixed with PBS + 2% FBS to 1 ml. The mixture was slowly added to the side of a SepMate tube (SepMate-15 IVD, Stem Cell Technologies) containing 4.5 ml of Lymphoprep (#07811, Stem Cell Technologies) and centrifuged for 20 minutes at 800 RPM. Approximately 900 μl of CPBMC layer was extracted below the plasma layer. To further remove red blood cells, 100 μl of red blood cell lysis buffer (eBioscience 10X RBC Lysis Buffer, #00-4300-54) was added to the CPBMCs and incubated at RT for 15 minutes. The CPBMC pellet was washed twice with PBS and centrifuged at 400 rpm for 5 minutes. Cells were counted using trypan blue via an automated detector (Countess II Automated Cell Counter) and subjects’ cells were pooled together for subsequent single-cell RNA sequencing.

### Single-cell RNA sequencing

Subject pooled single-cell suspensions were loaded onto a Chromium Single Cell Chip (10X Genomics) based on manufacturer’s instructions (targeted 10,000 cells per sample, 2,500 cells per person per time point). Captured mRNA was barcoded during cDNA synthesis and pooled for Illumina sequencing (Chromium Single Cell 3’ solution - 10X Genomics). Each time point was barcoded with a unique Illumina sample index, and then pooled together for sequencing in a single Illumina flow cell. The libraries were sequenced with an 8-base index read, 26-base read 1 containing cell-identifying barcodes and unique molecular identifiers (UMIs), and a 91-base read 2 containing transcript sequences on a NovaSeq 6000.

### Single-cell Dataset Generation

FASTQ files from Illumina were demultiplexed and aligned using Cell Ranger v3.0 (https://support.10xgenomics.com/single-cell-gene-expression/software/pipelines/latest/what-is-cell-ranger) and the hg19 reference genome with all options set to their defaults.

### Sample Demultiplexing

Subjects provided their raw data file from 23andMe (https://www.23andme.com/) sequencing for demultiplexing. Subjects S2 and S3 were genotyped via 23andMe’s V2 platform, and subjects S1 and S4 on 23andMe’s V4 platform. 23andMe files were then converted into variant call format (VCF) files using 2vcf (https://github.com/arrogantrobot/23andme2vcf/) against the hg19 (human) reference genome, which specifies where each subject’s genome differs from the human genome reference. VCF files were then indexed and sorted using vcftools (https://github.com/vcftools/vcftools). Demuxlet (https://github.com/statgen/demuxlet) was then run on each sample using the VCF files for each subject; this generates a probability of whether each cell barcode belongs to each subject, given the detection of single nucleotide polymorphism (SNPs) in reads associated with that cell barcode. Each cell was then assigned to the subject with the highest probability. Cells with low confidence (ambiguous cells) and high confidence in more than one subject (multiplets) were discarded, using demuxlet’s default confidence thresholds. See the README at https://github.com/thomsonlab/capblood-seq for detailed instructions.

### Debris Removal

The raw cell gene matrix provided by Cell Ranger contains gene counts for all barcodes present in the data. To remove barcodes representing empty or debris-containing droplets, a debris removal step was performed. First, a UMI count threshold was determined that yielded more than the expected number of cells based on original cell counts (15,000). All barcodes below this threshold were discarded. For the remaining barcodes, principal component analysis (PCA) was performed on the log-transformed cell gene matrix, and agglomerative clustering was used to cluster the cells. The number of clusters was automatically determined by minimizing the silhouette score among a range of numbers of clusters (6 to 15). For each cluster, a barcode dropoff trace was calculated by determining the number of barcodes remaining in the cluster for all thresholds in increments of 50. These cluster traces were then clustered into two clusters using agglomerative clustering - the two clusters representing “debris” with high barcode dropoff rates and “cells” with low barcode drop-off rates. All clusters categorized as “debris” were then removed from the data.

### Gene Filtering

Before cell typing, genes that have a maximum count less than 3 are discarded. Furthermore, after cell typing, any genes that are not present in at least 10% of one or more cell types are discarded.

### Data Normalization

Gene counts were normalized by dividing the number of times a particular gene appears in a cell (gene cell count) by the total gene counts in that cell. Furthermore, for visualization only, the gene counts were multiplied by a constant factor (5000), and a constant value of 1 was added to avoid zeros and then log transformed.

### Cell Typing

Dimensionality reduction was applied (PCA, number of components = 50) on the log-transformed gene expression cell matrix. The data was then subjected to t-distributed stochastic neighbor embedding (t-SNE) to project cells to 2 dimensions for manual annotation. Visually separable clusters were annotated based on known immune cell type markers (Fig. S1).

### Diurnal Gene Detection

To identify genes that exhibit diurnal variation in distinct cell types, we developed a statistical procedure that detects robust gene expression differences between morning (AM) and evening (PM) samples. Given that gene expression is different between subjects, we first normalize the mean gene expression within each subject for each cell type.

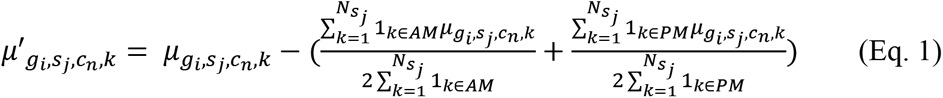

We take the mean gene expression *μ* for each gene *g_i_* in all samples *k* for cell type *c_n_* and subject *s_j_* and renormalize it into *μ*’ by subtracting the equally weighted mean of AM and PM samples (Eq. 1). We then split the mean gene values into an AM group and a PM group and perform a statistical test (two-tailed student-t test) to determine whether to reject the null hypothesis that gene expression in AM and PM samples come from the same distribution. We then perform Benjamini-Hochberg multiple comparison correction at an FDR of 0.05 on all gene and cell type p-values to determine where to plot the significance threshold. For plotting the genes, we choose the Z-statistic corresponding to the minimum p-value among cell types for that gene. To determine diurnality at the population level, we repeated the procedure above with all cells pooled into a single cell type.

### Subject and Cell Type Specific Gene Detection

To classify genes as subject specific, we detect genes with mean gene expression levels that are robustly different between subjects in at least one cell type. For each cell type *c_n_* and gene *g_i_*, we create subject groups containing the mean gene expression values from each sample. To determine whether the gene expression means from the different subjects do not originate from the same distribution, we perform an ANOVA one-way test to get an F-statistic and p-value for each gene. We then perform Benjamini-Hochberg multiple comparison correction at an FDR of 0.05 on all gene and cell type p-values. For plotting the genes, we chose the F-statistic corresponding to the minimum p-value among cell types for that gene.

For determining gene cell type specificity, we performed a similar procedure. In particular, for each gene *g_i_*, we create cell type groups containing the mean gene expression values for that cell type from each sample. We then perform a one-way ANOVA, and Benjamini-Hochberg multiple comparison correction at an FDR of 0.05.

### Pathway Enrichment Analysis

Pathways from the KEGG database (python bioservices package) were used to calculate pathway enrichment for genes that were among the top 250 most diurnal and individual specific. All remaining genes present in the data were considered background. In order to normalize for gene presence across pathways, each gene was weighted by dividing the number of pathways in which that gene appears. For each KEGG pathway, the test statistic for a two-proportion z-test (python statsmodel v0.11.1) is used to determine pathway enrichment. From the top level pathway classes, we broke out “Diseases” into “Other”, “Immune Diseases”, and “Infectious Diseases” and separated “Immune System” from “Organismal System” to understand diurnal and subject-specific genes in an immune relevant context.

## Supplementary Figures

**Supplementary Fig. 1.**
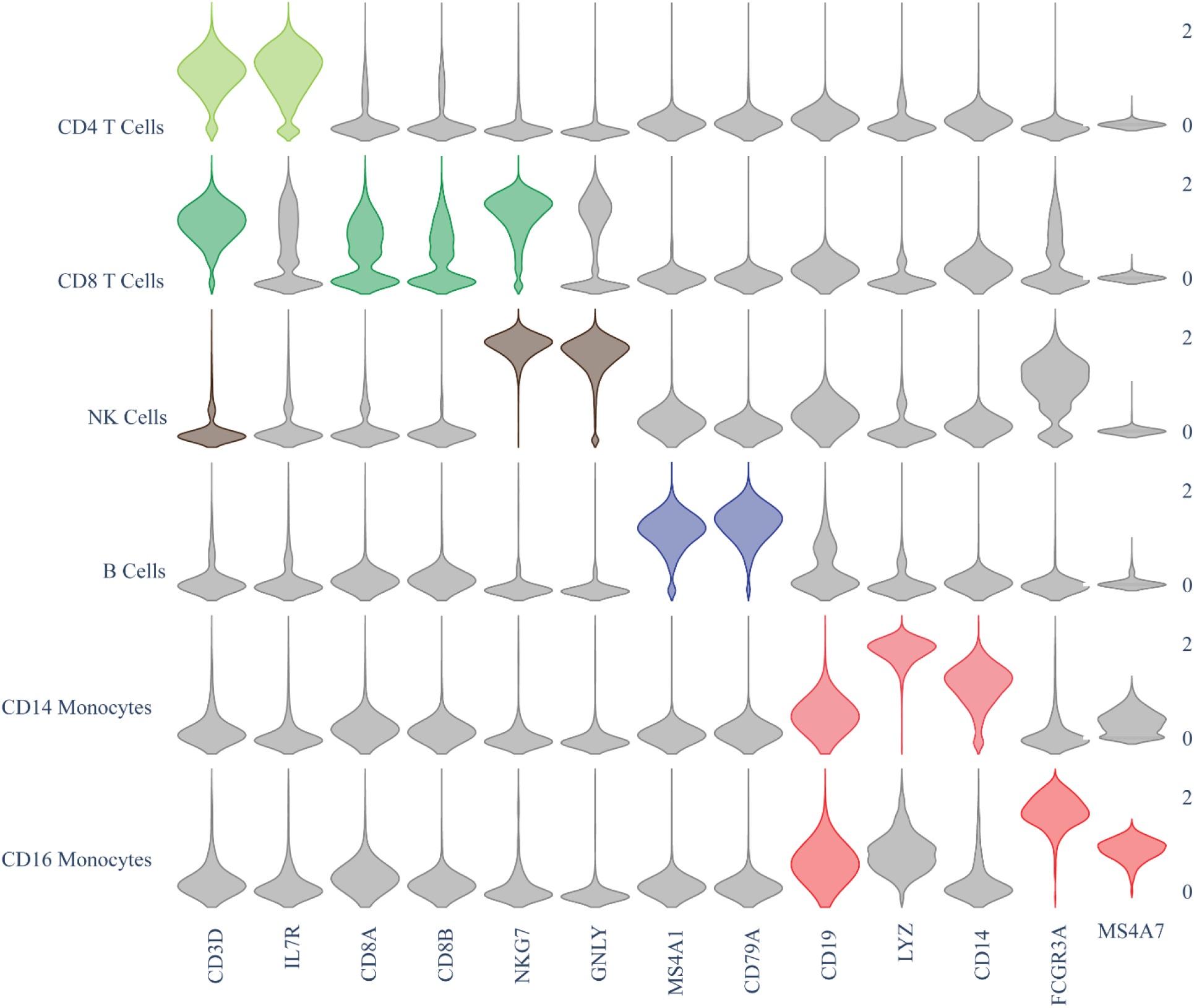
Cell type marker gene expression in cell clusters. Violin plots of log-normalized gene expression (y-axis, right hand side) for cell type markers (y-axis, left hand side) used to annotate cell clusters (x-axis) for known cell types. The colors correlate to clusters from Figure 1.d.

**Supplementary Fig. 2.**
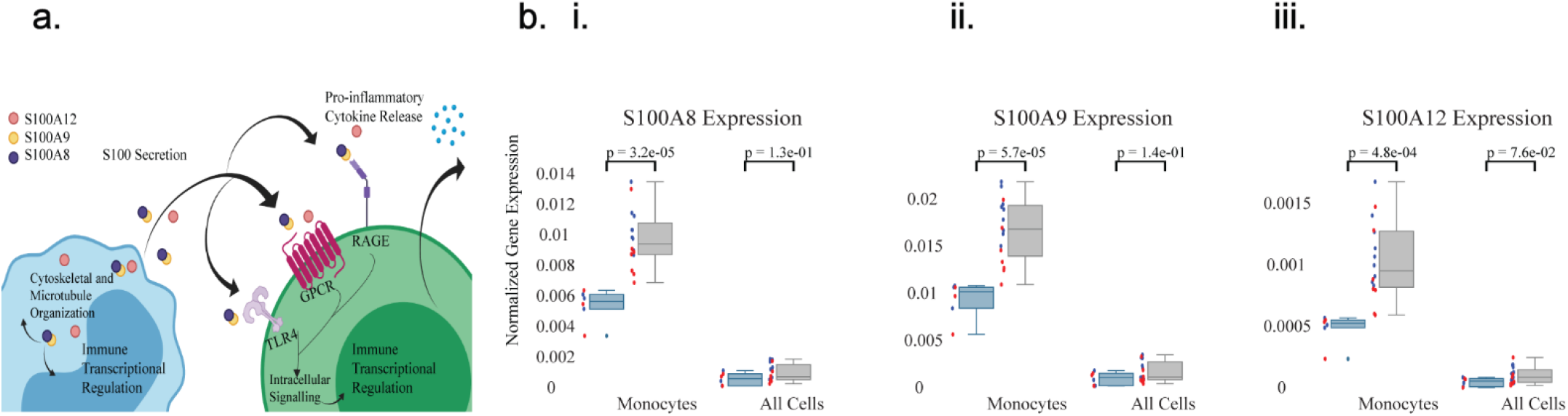
S100 pathway exhibits individual-specific regulation. **(a)** Simple schematic illustrating the role of S100A8, S100A9, and S100A12 genes in immune regulation. **(b)** Normalized mean gene expression of S100A8, S100A9, and S100A12 genes for S2 showing significant downregulation in monocytes as compared to all cells.

**Table S1:** Genes that ranked in top 20 that had pre-existing literature tying to circadian/diurnal expression

| Gene | DOI Reference |
| --- | --- |
| DDIT4 | 10.7554/eLife.20214.001,<br>10.1073/pnas.1800314115 |
| SMAP2 | 10.1038/s41398-019-0671-7 |
| RPL19 | 10.1128/MCB.00701-15 |
| RPS9 | 10.1073/pnas.1515308112 |
| PCPB1 | 10.1038/s41556-019-0441-z |
| RPS2 | 10.1073/pnas.1601895113 |
| RPL11 | 10.1073/pnas.252784499 |
| RBM3 | 10.1038/srep02054 |
| COX5B | 10.1152/physiolgenomics.00066.2007 |
| PDIA3 | 10.1002/jbmr.3046 |

